# Protocol to isolate oligodendrocytes, microglia, endothelial cells, astrocytes, and neurons from a single mouse brain using magnetic-activated cell sorting

**DOI:** 10.1101/2025.08.08.666877

**Authors:** Samah Houmam, Dominika Siodlak, Kevin D. Pham, Casandra Salinas-Salinas, Sarah R. Ocañas, Willard M. Freeman, Heather C. Rice

## Abstract

The isolation of specific cell types of the brain is essential to study cell-type-specific differences in complex neurological diseases such as Alzheimer’s disease. This protocol isolates oligodendrocytes, microglia, endothelial cells, astrocytes, and neurons from a single mouse brain. The process involves gentle tissue homogenization, debris removal, and sequential sorting of five distinct cell types. We validate cell purity and viability using flow cytometry and RT-qPCR. This protocol is well-suited for a range of downstream applications, including genomics, transcriptomics, and proteomics.

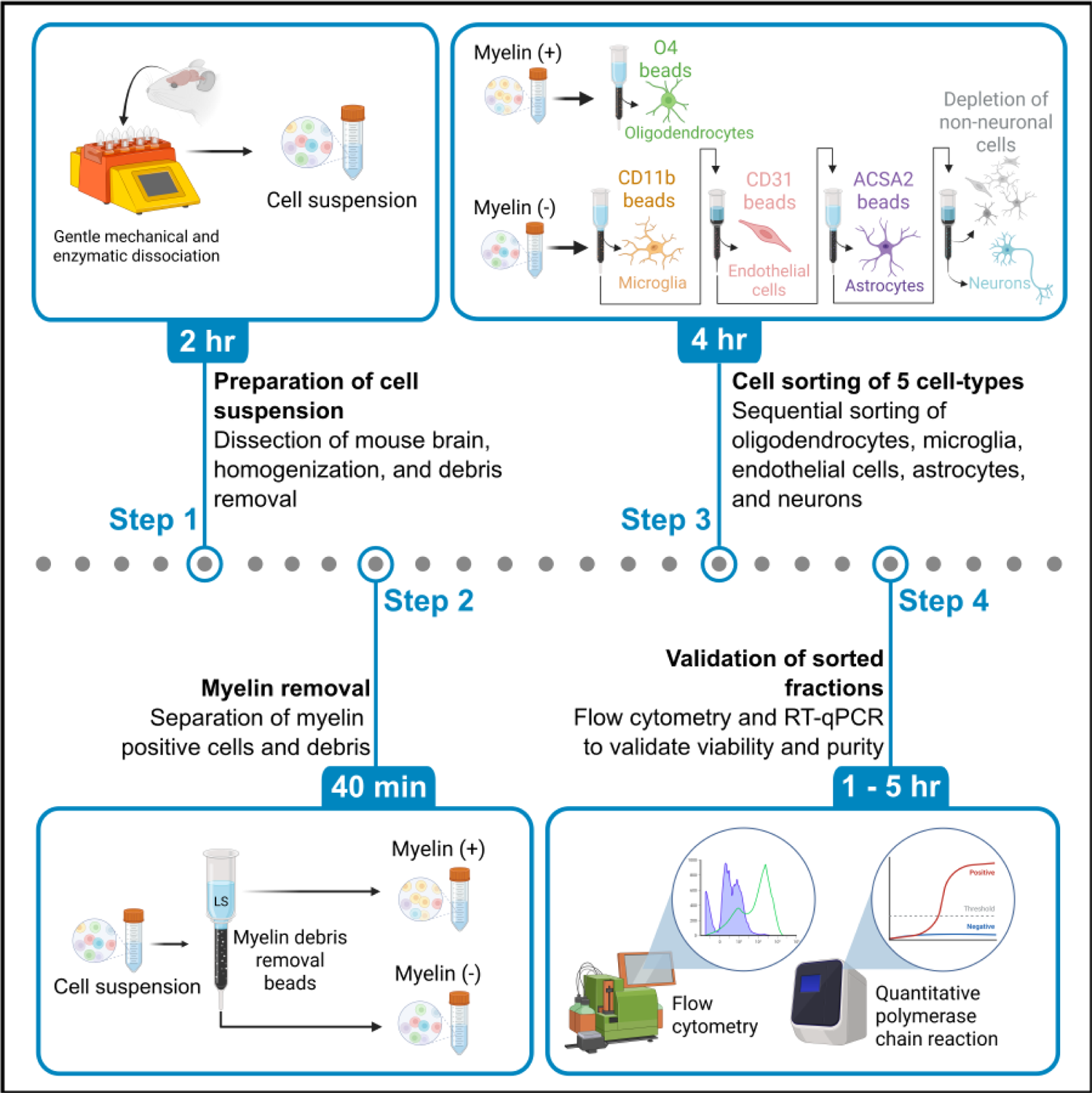

## Before you begin

The brain is a complex organ composed of diverse cell types, each contributing to the maintenance of homeostasis^1^. Isolating these distinct cell types is essential to distinguish their individual contributions to health and disease. While single-cell transcriptomics is a powerful tool to investigate transcriptional changes in individual cells, it has limitations such as reduced sensitivity for low-abundance transcripts and a 3’ bias that hampers detection of splice variants especially at the 5’ end^2^. In addition, single-cell transcriptomics can be expensive and select against fragile cell types underrepresenting them in the final clusters analyzed^3^. These limitations can be addressed by optimization of cell preparation conditions and bulk sorting cells of interest from the brain. Here, we present a protocol that uses gentle and rapid magnetic-activated cell sorting (MACS) to isolate five different cell types from a single mouse brain. This method is cost-effective, scalable, and compatible with parallel processing of multiple samples, increasing throughput. This protocol can be used for a variety of downstream applications such as studying the response of specific cell types to disease conditions, aging, or other stressors. Moreover, it can also be modified to suit various experimental needs, such as the sorting of a distinct cellular subtype, the use of specific brain regions, and different age time-points, making it a valuable protocol for the neuroscience research community.

## Innovation

Previous protocols using bulk sorting have not isolated all five cell types simultaneously ^4–6^, typically omitting either endothelial cells ^4,5^ or neurons ^6^. Additionally, they typically lack an assessment of endothelial cell contamination ^4,5^ which we observed to be present in the astrocytic and neuronal fractions without endothelial cell isolation prior to these steps. In addition, these previous reports typically lack consideration for the sensitivity of neurons to harsh sorting conditions and the risk for *ex vivo* activation of cells which was shown to occur in microglia^7^.To address these limitations, this protocol expands upon these prior studies^4–6^ by incorporating gentle enzymatic and mechanical dissociation with a trehalose-enhanced buffer to protect neurons^8^, and transcription and translation inhibitors to minimize *ex vivo* activation^7^. This is followed by sequential sorting of oligodendrocytes, microglia, endothelial cells, astrocytes, and neurons by MACS. The sorted cells can be used in a variety of downstream applications such as genomics, transcriptomics, and proteomics^9,10^ and can also be used to complement and enhance single-cell transcriptomic data^11^.

## Institutional permissions

All animal procedures described in this protocol were approved by the Institutional Animal Care and Use Committee of the Oklahoma Medical Research Foundation and performed in accordance with the National Institutes of Health (NIH) Guide for the Care and Use of Laboratory Animals. Users will need to acquire permission from the relevant institutions to perform animal work.

## Preparation of dissection and homogenization solutions

### Timing: 1h

**Note:** All volumes below are for one mouse brain. Scale up accordingly if processing more than one brain.

1. Prepare supplemented EBSS solution according to the recipe under the “Materials” section.
2. Prepare Worthington Papain Dissociation Kit Solutions

a. Add 32mL of EBSS (vial 1) to the albumin ovomucoid inhibitor mixture (vial 4). Allow to dissolve.
b. Add 5mL of supplemented EBSS solution to the papain vial (vial 2). Mix gently.
c. Add 500uL of supplemented EBSS solution to DNase vial (vial 3). Mix gently. Transfer 250uL of this solution to the papain vial (vial 2) and store the rest on ice until needed (step 6b).
d. Prepare a resuspension buffer by mixing 2.7mL of EBSS (vial 1) with 300uL of albumin ovomucoid inhibitor solution (vial 4).
e. Equilibrate the albumin ovomucoid inhibitor mixture (vial 4), papain solution (vial 2), and resuspension buffer using Carbogen gas mixture by flowing Carbogen on the surface of the liquid until pH is around 7.4. Use the pH color chart provided with the Worthington kit to assess the color of the phenol red in solution. **Critical:** Do not bubble gas directly into any of the solutions as this can result in frothing and compromise the integrity of contained proteins.
f. Transfer the papain solution (vial 2) to a MACS C-tube and add 5uL each of Anisomycin (10mg/mL), Actinomycin-D (5mg/mL), and Triptolide (10mM) stock solutions (1:1000 dilution).

## Key resources table

| REAGENT or RESOURCE | SOURCE | IDENTIFIER |
| --- | --- | --- |
| **Antibodies** |  |  |
| CD11b Antibody, anti-mouse, REAfinity, Vioblue, Clone REA592 \| M1/70.15.11.5 | Miltenyi | 130-113-810 |
| O4 Antibody, anti-human/mouse/rat, REAfinity, PE, Clone REA576 \| O4 | Miltenyi | 130-117-823 |
| CD34 Antibody, anti-mouse, REAfinity, FITC, Clone REA383 \| RAM34 | Miltenyi | 130-117-775 |
| ACSA-2 Antibody, anti-mouse, REAfinity, APC, Clone REA969 \| IH3-18A3 | Miltenyi | 130-116-245 |
| **Chemicals, peptides, and recombinant proteins** |  |  |
| DL-2-Amino-5-phosphonopentanoic acid (DL-AP5) | Millipore sigma | 165304-5MG |
| Kynurenic acid (KA) | Millipore sigma | K3375-1G |
| D-Trehalose | Millipore sigma | T9531-10G |
| Anisomycin | Millipore sigma | A9789-5MG |
| Actinomycin D | Millipore sigma | A1410-2MG |
| Triptolide | Millipore sigma | T3652-1MG |
| Dulbecco's phosphate buffered saline (DPBS) w/ calcium, magnesium, glucose, & pyruvate | Thermofisher | 14287080 |
| Bovine Serum Albumin (BSA) | Fisher | BP1600-100 |
| Dimethyl sulfoxide (DMSO) | Fisher | BP 231-100 |
| 7-AAD staining solution | Miltenyi | 130-111-568 |
| Trypan blue stain (0.4%) | Gibco | 15250-061 |
| **Critical commercial assays** |  |  |
| Worthington papain dissociation kit | Worthington | LK003150 |
| Myelin removal beads II, human, mouse, rat | Miltenyi | 130-096-733 |
| CD11b microbeads UltraPure, mouse | Miltenyi | 130-126-725 |
| Anti-ACSA-2 microbead kit, mouse | Miltenyi | 130-097-678 |
| CD31 microbeads, mouse | Miltenyi | 130-097-418 |
| Neuron isolation kit, mouse | Miltenyi | 130-115-389 |
| Anti-O4 microbeads, human, mouse, rat | Miltenyi | 130-096-670 |
| RNeasy kit | Qiagen | 74104 |
| Applied Biosystems High-Capacity cDNA Reverse Transcription Kit with RNase Inhibitor | Applied Biosystems | 4374966 |
| TaqMan™ Gene Expression Master Mix | Applied Biosystems | 4369016 |
| **Experimental models: Organisms/strains** |  |  |
| C57BL/6J | Jackson Laboratory | Strain #000664 |
| **Oligonucleotides** |  |  |
| Tmem119 TaqMan qPCR primer | Thermofisher | Mm00525305_m1 |
| Mbp TaqMan qPCR primer | Thermofisher | Mm01266402_m1 |
| Cldn5 TaqMan qPCR primer | Thermofisher | Mn00727012_s1 |
| Aldh1l1 TaqMan qPCR primer | Thermofisher | Mm03048957_m1 |
| Tubb3 TaqMan qPCR primer | Thermofisher | Mm01250301_g1 |
| β-actin TaqMan qPCR primer | Thermofisher | Mm02619580_g1 |
| **Software and algorithms** |  |  |
| GraphPad Prism 10 | GraphPad | www.graphpad.com |
| MACSQuantify™ 2.13 | Miltenyi | 130-094-556 |
| **Other** |  |  |
| GentleMACS™ C tubes | Miltenyi | 130-093-237 |
| LS columns | Miltenyi | 130-042-401 |
| LD columns | Miltenyi | 130-042-901 |
| MACS® SmartStrainers 70um | Miltenyi | 130-098-462 |
| QuadroMACS™ separator | Miltenyi | 130-090-976 |
| MACS® MultiStand | Miltenyi | 130-042-303 |
| Petri dish with lid | Fisher Scientific | FB0875713 |
| Decapitation scissors | Fisher Scientific | 28253 |
| Fine scissors | Fisher Scientific | 17-467-109 |
| Fine Science Tools Dumont #5 Forceps | Fisher Scientific | NC9889584 |
| Tapered spatula | Fisher Scientific | 2140110 |
| Weighing paper | Fisher Scientific | 09-898-12B |
| Razor blades, single-edged | Fisher Scientific | 12-640 |
| Absorbent disposable underpads | Carolina Biologicals | 567600A |
| 15mL Conical tube Sterile Polypropylene Centrifuge Tubes | Thermofisher | 339650 |
| 50mL Conical tube Sterile Polypropylene Centrifuge Tubes | Thermofisher | 339652 |
| Round-Bottom Polystyrene Test Tubes | Fisher Scientific | 14-959-1A |
| Pipette tips 10-1000uL | MidSci | PR-10XLRFL, PR-200RFL, 1250RFL |
| Serological Pipettes | Thermofisher | 170374N |
| Ice bucket | VWR | 10146-292 |
| gentleMACS™ Octo dissociator with heaters | Miltenyi | N/A |
| MACSQuant® analyzer 10 flow cytometer | Miltenyi | N/A |
| Sorvall Legend XTR centrifuge with TX-1000 rotor | Thermofisher | N/A |
| Eppendorf Centrifuge 5424 R | Eppendorf | N/A |
| BIORAD T100 Thermal Cycler | BIORAD | N/A |
| QuantStudio™ 5 | Applied Biosystems | N/A |

## Materials

### DL-AP5 stock solution

Materials DL-AP5 stock solution
| Reagent | Final concentration | Amount |
| --- | --- | --- |
| DL-AP5 | 100mM | 5mg |
| NaOH (1N) | n/a | 32.18uL |
| ddH<sub>2</sub>O | n/a | 289.7uL |

Prepare inside BSC. Aliquot and store at -20C.

### Kynurenic acid stock solution

| Reagent | Final concentration | Amount |
| --- | --- | --- |
| Kynurenic acid | 100mM | 11.35mg |
| NaOH (1N) | n/a | 60uL |
| ddH<sub>2</sub>O | n/a | 540uL |

Prepare inside BSC. Aliquot and store at -20C.

### Trehalose stock solution

| Reagent | Final concentration | Amount |
| --- | --- | --- |
| Trehalose | 50% | 10g |
| ddH<sub>2</sub>O | n/a | 20mL |

Autoclave, aliquot and store at -20C.

### Triptolide stock solution

| Reagent | Final concentration | Amount |
| --- | --- | --- |
| Triptolide | 10mM | 1mg |
| DMSO | n/a | 277.47uL |

Prepare inside BSC. Aliquot and store at -20C protected from light.
**CRITICAL:** This is a hazardous compound (acute oral and inhalation toxicity, reproductive toxin).
Please read the SDS carefully and handle it appropriately inside a BSC.

### Anisomycin stock solution

| Reagent | Final concentration | Amount |
| --- | --- | --- |
| Anisomycin | 10mg/mL | 5mg |
| DMSO | n/a | 500uL |

Prepare inside BSC. Aliquot and store at -20C protected from light.
**CRITICAL:** This is a hazardous compound (acute oral toxicity). Please read the SDS carefully and handle it appropriately inside a BSC.

### Actinomycin-D stock solution

| Reagent | Final concentration | Amount |
| --- | --- | --- |
| Actinomycin-D | 5mg/mL | 2mg |
| DMSO | n/a | 400uL |

Prepare inside BSC. Aliquot and store at -20C protected from light.
**CRITICAL:** This is a hazardous compound (acute oral toxicity, carcinogen, reproductive toxin). Please read the SDS carefully and handle it appropriately inside a BSC.

### DPBS/BSA solution

| Reagent | Final concentration | Amount |
| --- | --- | --- |
| BSA | 0.5% | 2.5g |
| DPBS | n/a | 500mL |

Prepare inside BSC. Store at 4°C for up to a week.

### To be made fresh the day of the procedure

#### Supplemented EBSS

| Reagent | Final concentration | Amount |
| --- | --- | --- |
| EBSS (Worthington kit vial 1) | n/a | 5.34mL |
| AP5 (100mM) | 0.05mM | 3uL |
| KA (100mM) | 0.8mM | 48uL |
| Trehalose (50%) | 5% | 600uL |

Prepare inside laminar flow hood the day of the experiment. Store at room temperature until needed.
**Note:** These amounts are for one adult mouse brain. Scale accordingly if processing more than one brain.

### Equipment setup

1. At the beginning of the experiment day, turn on the flow cytometer and perform calibration/clean-up.
2. Prepare dissection area

a. On an absorbent pad, set out all dissection tools and a razor blade (**Figure 1A**)
b. In an ice bucket, place a 15mL conical tube filled with DPBS. This will be used to rinse the brain before chopping. Also, place a cold block topped with DPBS-wetted weighing paper in the ice bucket. This will be used as a surface for chopping the brain tissue (**Figure 1B**). **Note:** If a cold block is not available, an inverted petri dish can be used instead.
3. Prepare cell-sorting area

a. In a laminar flow hood, position an ice bucket under a QuadroMACS separator mounted on a MultiStand (**Figure 1C**). **Note:** The cell-sorting can also be performed in a biosafety cabinet (BSC) or on a laboratory bench.

## Step-by-step method details

### Dissection and dissociation of mouse brain tissue

#### Timing: 45min

1. Anesthetize the animal with isoflurane (3-5% induction). Verify depth of anesthesia using the toe-pinch method. Euthanize the mouse by cervical dislocation followed by decapitation. **Note:** Transcardiac perfusion with saline can be performed prior to tissue collection to eliminate blood-derived contamination. This step is particularly important for downstream applications such as RNA sequencing, proteomics, or flow cytometry, where residual blood cells and plasma proteins may confound experimental results.
2. Dissect the brain out and quickly submerge it in the 15mL conical tube of ice-cold DPBS to rinse.
3. Transfer the brain to the cold block topped with DPBS-wetted weighing paper. Dissect the brain region of interest. Chop tissue into 3-4mm wide pieces using a razor blade. **Note:** Example data presented herein were derived from the cerebrum after removing the cerebellum and olfactory bulbs.
4. Transfer the tissue into the C-tube containing the papain-DNase solution with transcription/translation inhibitors.
5. Tightly close and attach the C tube upside down on the gentleMACS Octo Dissociator with Heaters. Run the 37C_ABDK_01 program. **Note:** Use the 37C_ABDK_02 program if your dissected tissue weighs <100mg. Preparation of cell suspension **Timing: 1h10min**
6. Pellet the homogenate and resuspend in resuspension buffer.

a. After completion of the program, centrifuge the homogenized solution at 300g for 3min at room temperature.
b. During this spin add 150uL of the remaining DNase solution to the resuspension buffer (Preparation of dissection and homogenization solutions – step 2c).
c. Discard supernatant and immediately resuspend pellet in the resuspension buffer (3.15mL) by gently pipetting up and down.
7. Prepare discontinuous density gradient.

a. In a 15mL conical tube, pipette 5mL of albumin ovomucoid inhibitor then carefully layer the resuspended homogenate on top (**Figure 1D**). Use a pipette controller and tilt the tube for slow and controlled pipetting.
b. Centrifuge at 300g for 5min at room temperature.
c. During this spin, place a 70uM MACS SmartStrainer on a 50mL conical tube and pre-wet with 2mL of DPBS-BSA.
d. At the end of the spin, discard the supernatant (**Figure 1E**) and resuspend the pellet in 8mL of DPBS-BSA. Filter the cell suspension through the prepared strainer.
e. Perform a cell count using trypan blue staining to determine cell yield.
f. Take aliquots of filtered cell suspension to use as input in later analyses – 30uL for flow cytometry and 500uL for RT-qPCR is appropriate.
g. Centrifuge both cell suspension and the 500uL “input” aliquot to pellet the cells at 300g for 3min at 4°C. **Note:** All RT-qPCR pelleted aliquots will be lysed during the incubation with microbeads for the following isolation to save time. **Critical:** All incubations at 4°C should be performed in a refrigerator and not on ice.
8. Lyse the pelleted “input” fraction in 350uL of RLT Plus lysis buffer containing β-mercaptoethanol (provided with Qiagen RNEasy kit). Store in the −80°C freezer until ready to isolate RNA. **Note:** To save time, this can be done during the incubation of step 10. Myelin removal **Timing: 40min**
9. Resuspend the cell pellet (from step 7g) in 150uL of DPBS-BSA and add 30uL of myelin removal beads. Incubate at 4°C in the dark for 10 min.
10. During this incubation, mount an LS column on the QuadroMACS stand with its wings facing forward (**Figure 1F**). Position a 15mL conical tube in a cold rack or ice bucket under the column. Pipette 2mL of DPBS-BSA onto the column and let it run through the column to pre-wet it. **Note:** During this incubation, step 8 can be performed.
11. Wash the cells by adding 1mL of DPBS-BSA to the conical tube (step 9) and centrifuge at 300g for 3min at 4°C.
12. Discard the supernatant. Resuspend the cell pellet in 500uL of DPBS-BSA and load onto the column. Collect the flow-through.
13. Wash by adding 3mL of DPBS-BSA onto the column and collect the flow-through. When flow stops, repeat with another 3mL of DPBS-BSA and continue collecting the flow-through.
14. To eject the myelin-positive fraction, remove the column from the stand and place it on a labeled 15mL conical tube. Then, add 5mL of DPBS-BSA to the column and quickly use the plunger to flush out the magnetically labeled cells. Firmly and consistently push the plunger into the column. There will be foaming at the end, this is normal.
15. Centrifuge the myelin fraction (column-ejected) and flow-through (myelin-negative) at 300g for 3min at 4°C to pellet the cells. The myelin fraction will be used to isolate oligodendrocytes (steps 16-25), and the flowthrough will be used to isolate microglia, endothelial cells, astrocytes, and neurons (steps 26-62). Isolation of oligodendrocytes **Timing: 45min** **Note:** This step can be performed side by side with the following microglial isolation step.
16. Resuspend the myelin fraction cell pellet (from step 15) in 150uL of DPBS-BSA and add 30uL of anti-O4 microbeads. Incubate at 4°C in the dark for 10 min.
17. During this incubation, mount an LS column on the QuadroMACS stand with its wings facing forward (**Figure 1F**). Position a 15mL conical tube in a cold rack or ice bucket under the column. Pipette 2mL of DPBS-BSA onto the column and let it run through the column to pre-wet it.
18. At the end of the incubation, wash the cells by adding 1mL of DPBS-BSA to the conical tube (step 16) and centrifuge at 300g for 3min at 4°C.
19. Discard the supernatant. Resuspend the pellet in 500uL of DPBS-BSA and load onto the column and collect the flow-through.
20. Wash by adding 3mL of DPBS-BSA onto the column and collect the flow-through. When the flow stops, repeat with another 3mL and continue collecting the flow-through.
21. To eject the oligodendrocytes, remove the column from the stand and place it on a labeled 15mL conical tube. Add 5mL of DPBS-BSA to the column and quickly use the plunger to flush out the magnetically labeled cells. Firmly and consistently push the plunger into the column. There will be foaming at the end, this is normal. Set aside a 30uL aliquot for flow cytometry analysis.
22. Perform a cell count using trypan blue staining and a hemocytometer to determine cell yield.
23. Centrifuge the oligodendrocyte fraction (column-ejected) at 300g for 3min at 4°C to pellet the cells.
24. Discard the column flow-through.
25. Lyse the pelleted oligodendrocyte fraction in 350uL of RLT Plus lysis buffer containing β-mercaptoethanol (provided with Qiagen RNEasy kit). Store in the −80°C freezer until ready to isolate RNA. **Note:** To save time, this can be done during the incubation of step 36. Isolation of microglia **Timing: 45min** **Note:** This step can be performed side by side with the previous oligodendrocyte isolation step.
26. Resuspend the cell pellet from the flow-through (step 15; myelin-negative) in 150uL of DPBS-BSA and add 30uL of anti-CD11b beads. Incubate at 4°C in the dark for 10 min.
27. During this incubation, mount an LS column on the QuadroMACS stand with its wings facing forward (**Figure 1F**). Position a 15mL conical tube in a cold rack or ice bucket under the column. Pipette 2mL of DPBS-BSA onto the column and let it run through the column to pre-wet it.
28. At the end of the incubation, wash the cells by adding 1mL of DPBS-BSA to the conical tube (step 26) and centrifuge at 300g for 3min at 4°C.
29. Discard the supernatant. Resuspend the pellet in 500uL of DPBS-BSA and load onto the column and collect the flow-through.
30. Wash by adding 3mL of DPBS-BSA onto the column and collect the flow-through. When the flow stops, repeat with another 3mL and continue collecting the flow-through.
31. To eject the microglia, remove the column from the stand and place on a labeled 15mL conical tube. Add 5mL of DPBS-BSA to the column and quickly use the plunger to flush out the magnetically labeled cells. Firmly and consistently push the plunger into the column. There will be foaming at the end, this is normal. Set aside a 375uL aliquot for flow cytometry analysis.
32. Perform a cell count using trypan blue staining and a hemocytometer to determine cell yield.
33. Centrifuge the flow-through (rest of the cells) and microglial fraction (column-ejected) at 300g for 3min at 4°C to pellet the cells.
34. Lyse the pelleted microglial fraction in 350uL of RLT Plus lysis buffer containing β-mercaptoethanol (provided with Qiagen RNEasy kit). Store in the −80°C freezer until ready to isolate RNA. **Note:** To save time, this can be done during the incubation of step 36. Isolation of endothelial cells **Timing: 1hr**
35. Resuspend the cell pellet from the flow-through (step 33) in 150uL of DPBS-BSA and add 30uL of anti-CD31 microbeads. Incubate at 4°C in the dark for 10 min.
36. During this incubation, mount an LD column on the QuadroMACS stand with its wings facing forward (**Figure 1F**). Position a 15mL conical tube in a cold rack or ice bucket under the column. Pipette 2mL of DPBS-BSA onto the column and let it run through the column to pre-wet it. **Critical:** Do not use an LS column during this step. **Note:** During this incubation, steps 25 & 34 can be performed.
37. At the end of the incubation, wash the cells by adding 1mL of DPBS-BSA to the conical tube (step 35) and centrifuge at 300g for 3min at 4°C.
38. Discard the supernatant. Resuspend the pellet in 500uL of DPBS-BSA and load onto the column and collect flow-through.
39. Wash by adding 1mL of DPBS-BSA onto the column and collect the flow-through. Repeat with another 1mL when flow stops and keep collecting flow-through. **Note:** LD columns have a much slower flow rate than LS columns
40. To eject the endothelial cells, remove the column from the stand and place on a labeled 15mL conical tube. Add 3mL of DPBS-BSA to the column and quickly use the plunger to flush out the magnetically labeled cells. Firmly and consistently push the plunger into the column. There will be foaming at the end, this is normal. Set aside a 225uL aliquot for flow cytometry analysis.
41. Perform a cell count using trypan blue staining and a hemocytometer to determine cell yield.
42. Centrifuge the flow-through (rest of the cells) and endothelial fraction (column-ejected) at 300g for 3min at 4°C to pellet the cells.
43. Lyse the pelleted endothelial fraction in 350uL of RLT Plus lysis buffer containing β-mercaptoethanol (provided with Qiagen RNEasy kit). Store in the −80°C freezer until ready to isolate RNA. **Note:** To save time, this can be done during the incubation of step 46. Isolation of astrocytes **Timing: 45min**
44. Resuspend the cell pellet from the flow-through (step 42) in 150uL of DPBS-BSA and add 30uL of FcR Blocking Reagent. Incubate at 4°C in the dark for 10 min.
45. Add 30uL of Anti-ACSA-2 microbeads to the same suspension and incubate at 4°C in the dark for another 10 min.
46. During this incubation, mount an LS column on the QuadroMACS stand with its wings facing forward (**Figure 1F**). Position a 15mL conical tube in a cold rack or ice bucket under the column. Pipette 2mL of DPBS-BSA onto the column and let it run through the column to pre-wet it. **Note:** During this incubation, step 43 can be performed.
47. At the end of the incubation, wash the cells by adding 1mL of DPBS-BSA to the conical tube (step 45) and centrifuge at 300g for 3min at 4°C.
48. Discard the supernatant. Resuspend the pellet in 500uL of DPBS-BSA and load onto the column and collect the flow-through.
49. Wash by adding 3mL of DPBS-BSA onto the column and collect the flow-through. When the flow stops, repeat with another 3mL and continue collecting the flow-through.
50. To eject astrocytes, remove the column from the stand and place on labeled 15mL conical tube. Add 5mL of DPBS-BSA to the column and quickly use the plunger to flush out the magnetically labeled cells. Firmly and consistently push the plunger into the column. There will be foaming at the end, this is normal. Set aside a 375uL aliquot for flow cytometry analysis.
51. Perform a cell count using trypan blue staining and a hemocytometer to determine cell yield.
52. Centrifuge the flow-through (rest of the cells) and astrocyte fraction (column-ejected) at 300g for 3min at 4°C to pellet the cells.
53. Lyse the pelleted astrocyte fraction in 350uL of RLT Plus lysis buffer containing β-mercaptoethanol (provided with Qiagen RNEasy kit). Store in the −80°C freezer until ready to isolate RNA. **Note:** To save time, this can be done during the incubation of step 56. Isolation of neurons **Timing: 50min**
54. Resuspend the cell pellet from the flow-through (step 52) in 150uL of DPBS-BSA and add 30uL of non-neuronal cell biotin-antibody cocktail. Incubate at 4°C in the dark for 10 min.
55. Add 30uL of Anti-biotin microbeads to the same suspension and incubate at 4°C in the dark for another 10 min.
56. During this incubation, mount an LD column on the QuadroMACS stand with its wings facing forward (**Figure 1F**). Position a 15mL conical tube in a cold rack or ice bucket under the column. Pipette 2mL of DPBS-BSA onto the column and let it run through the column to pre-wet it. **Note:** LD columns flow considerably slower than LS columns. **Note:** During this incubation, step 53 can be performed.
57. At the end of the incubation, wash the cells by adding 1mL of DPBS-BSA to the conical tube (step 55) and centrifuge at 300g for 3min at 4°C.
58. Discard the supernatant. Resuspend the pellet in 500uL of DPBS-BSA and load onto the column and collect the flow-through.
59. Wash by adding 1mL of DPBS-BSA onto the column and collect the flow-through. When the flow stops, repeat with another 1mL and continue collecting the flow-through.
60. Discard the column (rest of cells) and centrifuge the flow-through (neurons) at 300g for 3min at 4°C to pellet the cells. Set aside a 225uL aliquot for flow cytometry analysis.
61. Perform a cell count using trypan blue staining and a hemocytometer to determine cell yield.
62. Lyse pelleted neuronal fraction in 350uL of RLT Plus lysis buffer containing β-mercaptoethanol (provided with Qiagen RNEasy kit). Store in the −80°C freezer until ready to isolate RNA. Validation of fraction purity **Timing: variable RT-qPCR**
63. Extract RNA from all lysed fraction samples using the Qiagen RNEasy kit and quantify using a nanodrop instrument.
64. Convert RNA samples to cDNA using the Applied Biosystems High-Capacity cDNA Reverse Transcription Kit with RNAse Inhibitor.
65. Perform qPCR to probe each sample for the chosen cell type-specific markers (**Figure 2**). See also “Quantification and data analysis” section. Flow cytometry
66. For each cell type, use a third of the aliquot that was set aside for flow cytometry to prepare a stained sample. Use another third for an unstained control. Save the last third for viability staining (step 72).
67. Use an antibody panel to target cell-surface markers of your choice or utilize the panel we provided in this protocol (See Key Resource Table – Antibodies section).
68. Add antibodies to stained samples at the recommended dilution ratio. If using the provided panel, 1:50.
69. Incubate stained samples for 10min at 4°C in the dark.
70. Wash by adding 1mL of DPBS-BSA to the samples and centrifuge at 300g for 3min at 4°C.
71. Resuspend the pellets in 125uL of DPBS-BSA and run on a flow cytometer using an appropriate analysis template.
72. For each cell type, add 7-AAD (at a 1:1000 dilution) to the last third of the set-aside aliquot. No wash is needed. **Critical:** Add the 7-AAD 5 min or less before running samples on the flow cytometer as the dye can become toxic to the cells at longer incubation times.
73. Analyze results using appropriate gating strategy (**Figure 3-5**). See also “Quantification and data analysis” section.

**Figure 1:**
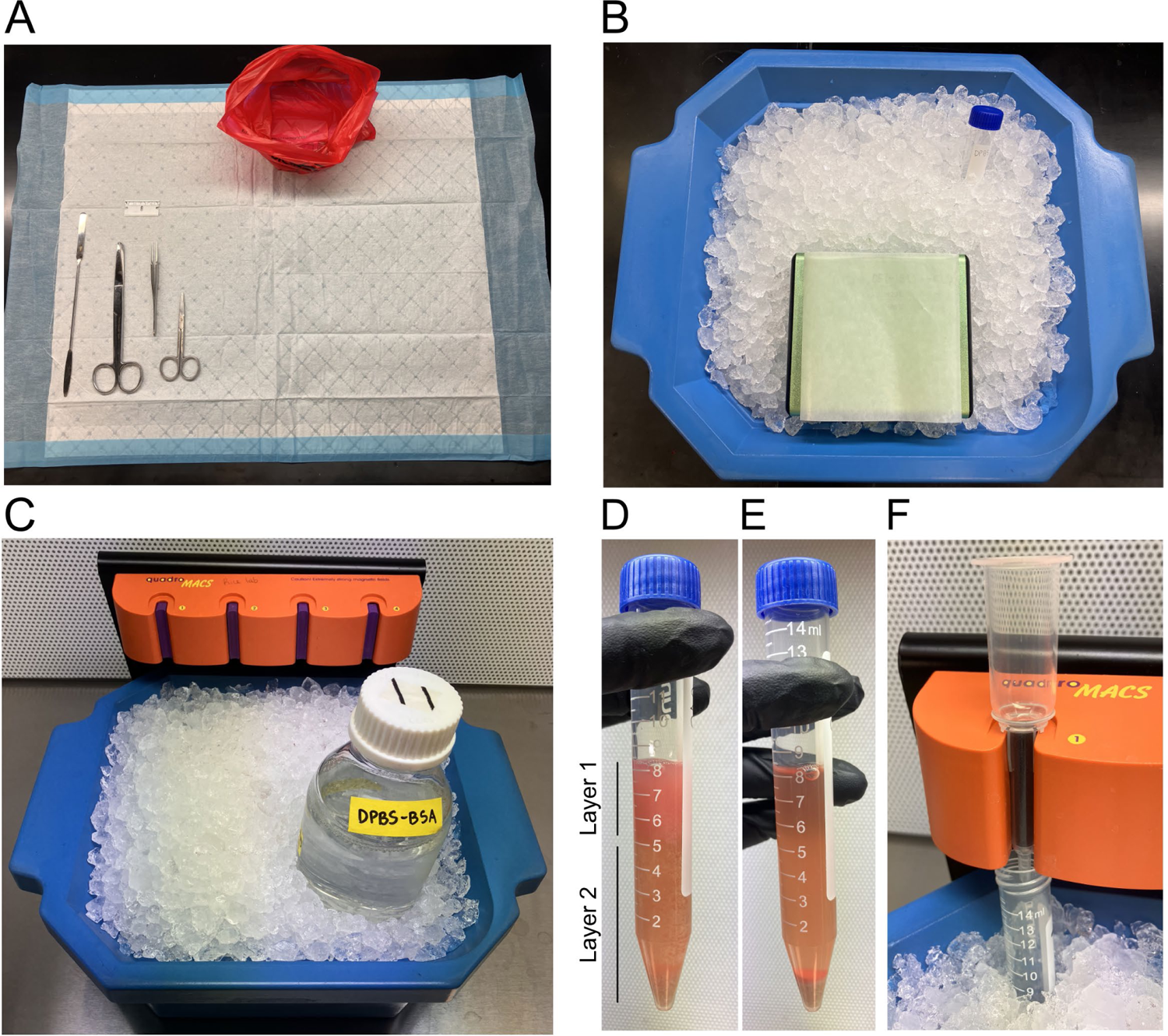
Set-up of dissection and cell-sorting area. (A) Dissection area and tools include decapitation scissors, fine scissors, fine forceps, tapered spatula, and a razor blade. (B) Cold surface set-up for cutting the dissected brain into smaller pieces. (C) Cell-sorting set up with the QuadroMACS stand and an ice bucket below to keep the flowthrough cold. Image of the debris removal gradient before (D) and after (E) centrifugation. (F) Separation column on QuadroMACS stand with wings facing forward.

**Figure 2:**
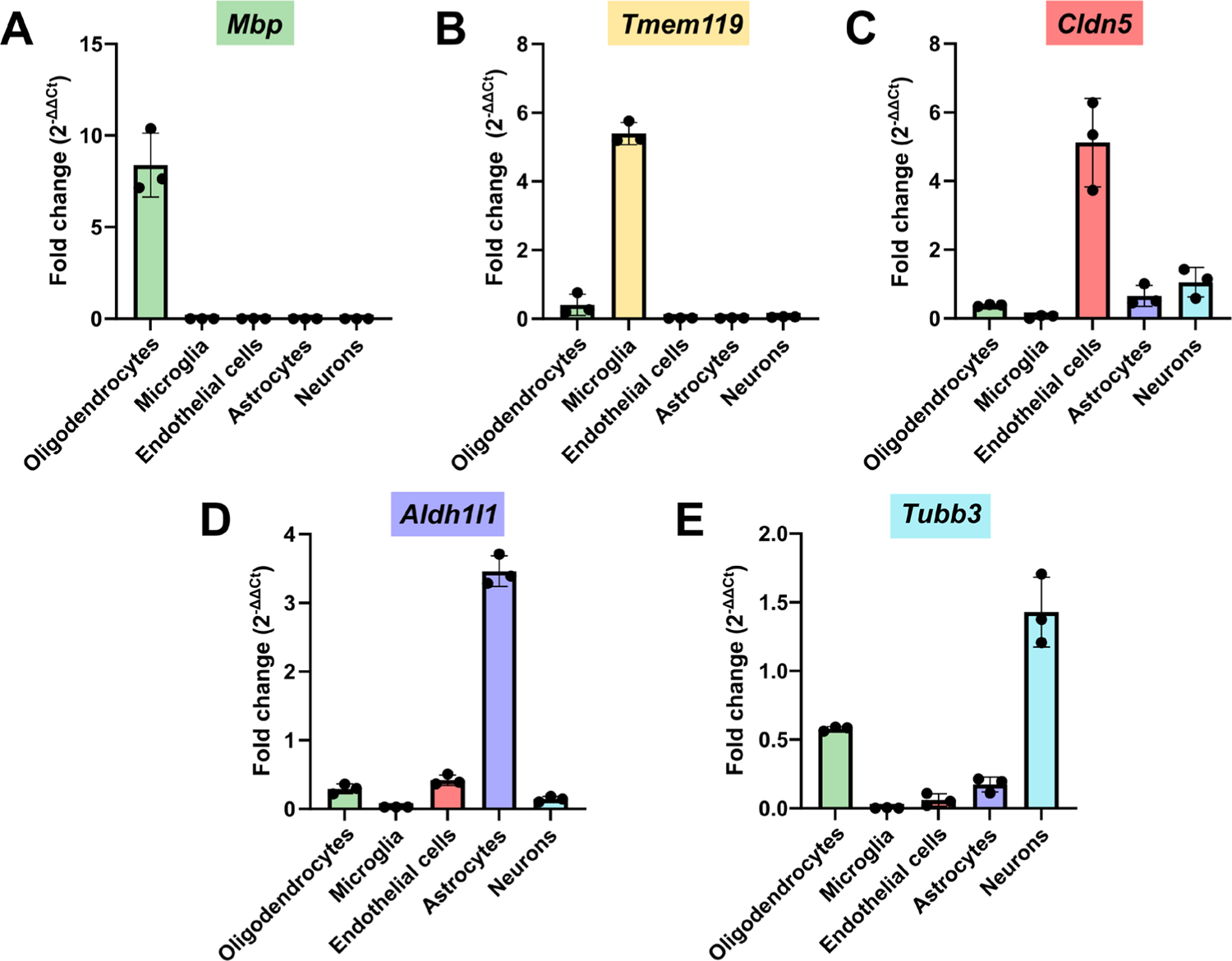
Quantitative RT-PCR validation of cell type-specific marker enrichment. RT-qPCR using cell-specific markers shows enrichment of (A) *Mbp* in the oligodendrocyte fraction, (B) *Tmem119* in the microglial fraction, (C) *Cldn5* in the endothelial fraction, (D) *Aldh1l1* in the astrocytic fraction, and (E) *Tubb3* in the neuronal fraction (n=3, mean ± SD).

**Figure 3:**
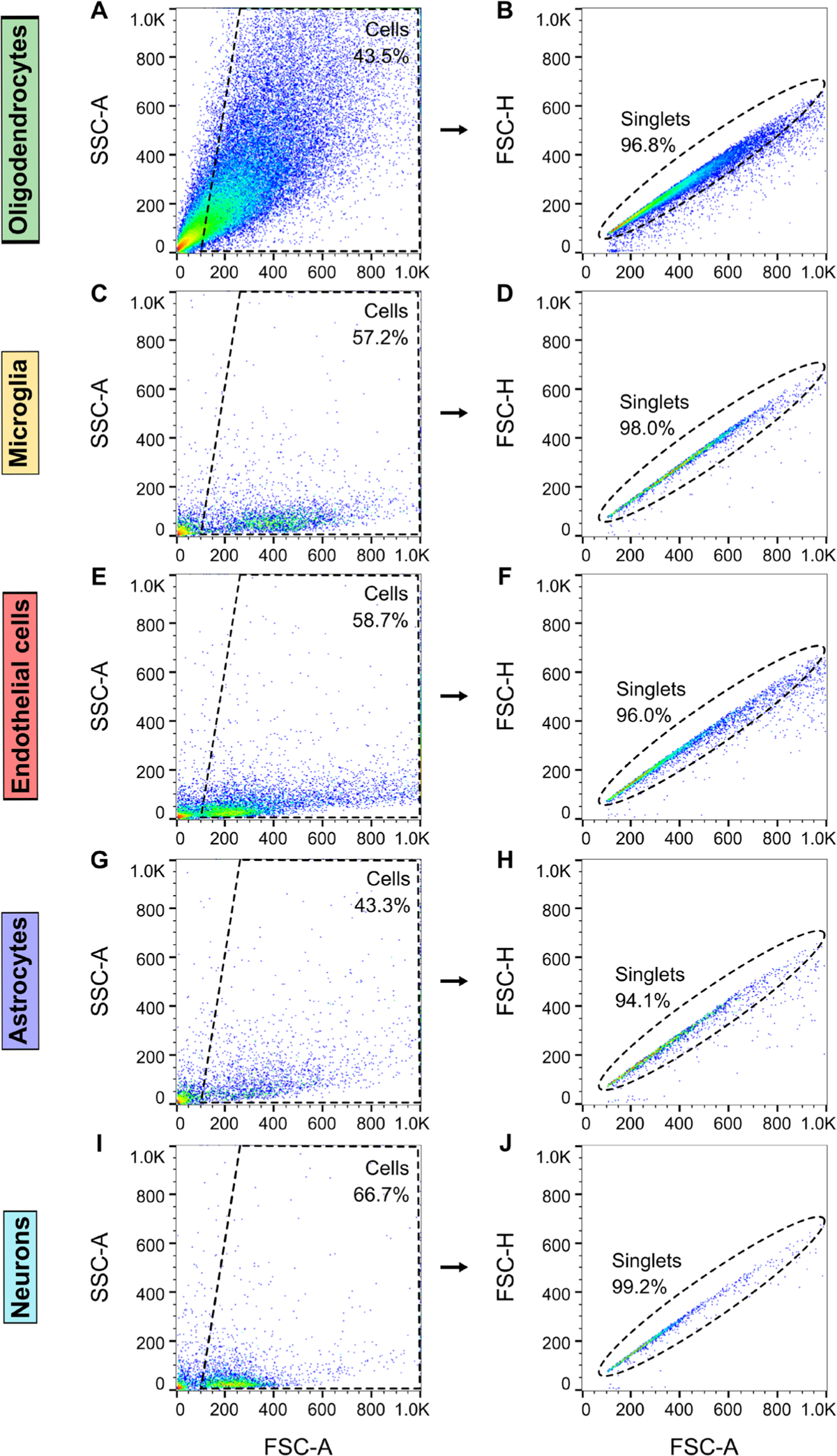
Flow-cytometry gating strategy. Representative scatter plots from a single experiment showing SSC-A versus FSC-A (A,C,E,G,I) to determine cell populations and exclude debris, and FSC-H versus FSC-A (B,D,F,H,J,L) to select singlets and exclude doublets for oligodendrocytes (A-B), microglia (C-D), endothelial cells (E-F), astrocytes (G-H), and neurons (I-J).

## Anticipated Results

Upon completion of this protocol, enriched cell populations of oligodendrocytes, microglia, endothelial cells, astrocytes, and neurons are isolated from a single mouse brain for subsequent analysis. Presented here are three independent replicates, each processed on a different day, which demonstrates consistency and reliability of the protocol. Each brain yields approximately 16 million oligodendrocytes, 1 million each of microglia, endothelial cells, and neurons, and 300,000 astrocytes, as determined by trypan blue staining of live cells and hemocytometer counting (**Table 1**). Cell viability, assessed by 7-AAD staining, exceeded 95% for oligodendrocytes, microglia, and neurons, and 70% for astrocytes and endothelial cells (**Table 2 and Figure 5**). To assess the purity of each cell fraction, we used a combination of RT-qPCR and flow cytometry. RT-qPCR analysis confirmed enrichment of the cell-type-specific markers within each respective cell fraction (**Figure 2**). Flow cytometry analysis demonstrates an enrichment from input of 99% for oligodendrocytes, 79% for microglia, 45% for endothelial cells, 74% for astrocytes, and >96% for neurons each in their respective isolated fraction, as indicated by the corresponding cell type-specific antibody signal (**Figure 4 and Table 3**). The endothelial cell fraction shows only a 45% enrichment of the CD34 marker using flow cytometry. While the endothelial cell fraction shows some staining with the O4 (oligodendrocyte marker) and ACSA2 (astrocyte marker) antibodies in flow cytometry (12-14%) suggesting potential contamination with oligodendrocytes and astrocytes respectively, RT-qPCR for the markers specific to oligodendrocytes (*Mbp)* and astrocytes *(Aldh1l1*) does not show enrichment in the endothelial fraction. Similarly, the astrocyte fraction shows some staining with the CD34 antibody (29%) which is not corroborated by the RT-qPCR data *(Cldn5)*. This suggests that the poor enrichment of the endothelial fraction and potential contamination of the astrocyte fraction as indicated by flow cytometry may not be due entirely to an impure fraction. Flow cytometry analysis of endothelial cells and astrocytes may require further optimization (See Troubleshooting section – Problem 1 and Limitations section). Lastly, astrocytes and endothelial cells show higher cell death levels during flow cytometry (∼20%. **Table 2**) which may affect their enrichment parameters as well.

**Figure 4:**
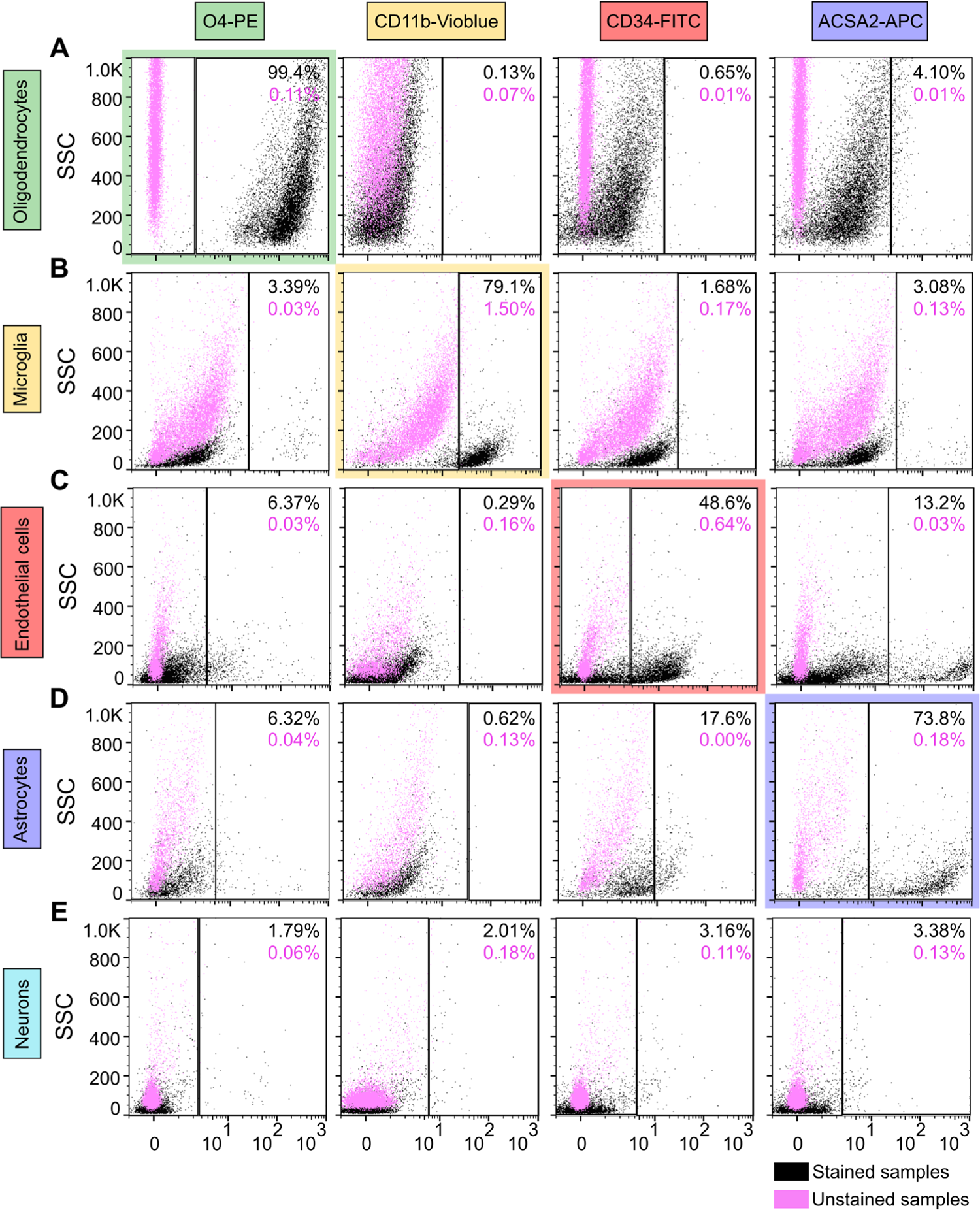
Establishment of flow cytometry gates using stained samples and unstained controls. Representative scatter plots from a single experiment showing stained samples (black) and unstained negative control samples (pink) for each cell-specific antibody (O4 as a marker for oligodendrocytes, CD11b for microglia, CD34 for endothelial cells, and ACSA2 for astrocytes) used to set the negative and positive staining gates for the (A) oligodendrocyte, (B) microglia, (C) endothelial cell, (D) astrocyte, and (E) neuron fractions.

**Figure 5:**
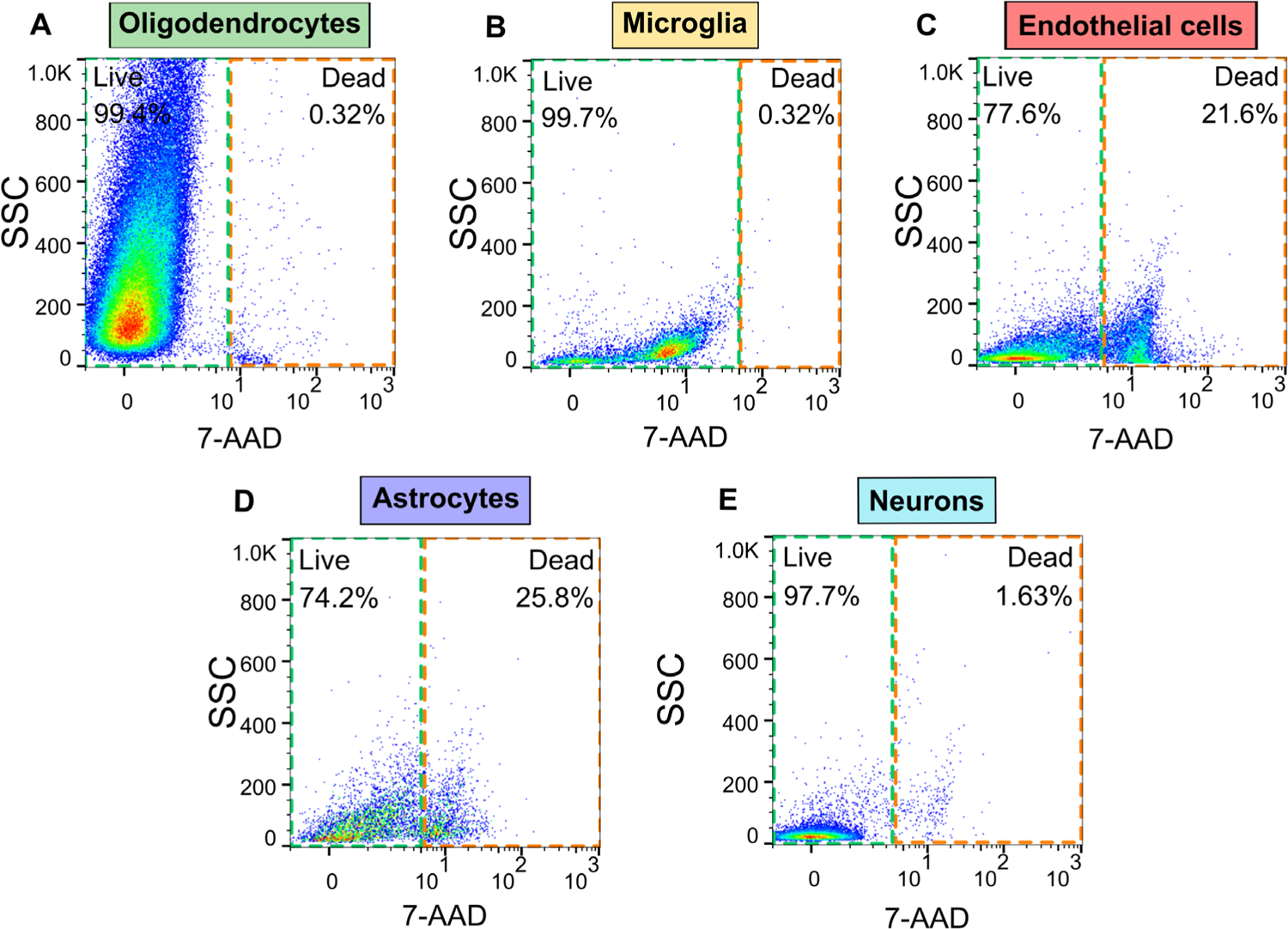
Viability assessment of cell-sorted fractions using 7-AAD staining. Representative scatter plots from a single experiment showing 7-AAD staining in purified cell fractions to determine live and dead populations in (A) oligodendrocytes, (B) microglia, (C) endothelial cells, (D) astrocytes, and (E) neurons.

**Table 1:** Summary of cell yield for each sorted cell type. Table of hemocytometer cell counts using Trypan blue staining (n=3 data displayed as average ± standard deviation).

| Cell type | Cell counts |
| --- | --- |
| Oligodendrocytes | 16mi ± 1mi |
| Microglia | 1mi ± 70K |
| Endothelial cells | 900K ± 200K |
| Astrocytes | 325K ± 40K |
| Neurons | 1mi ± 180K |

**Table 2:** Cell viability assessment by flow cytometry. Table summarizing percent live cells using 7-AAD staining (n=3, data displayed as average ± standard deviation).

| Cell type | Live cells (%) |
| --- | --- |
| Oligodendrocytes | 99.47 ± 0.09 |
| Microglia | 99.53 ± 0.12 |
| Endothelial cells | 72.70 ± 5.83 |
| Astrocytes | 71.90 ± 6.61 |
| Neurons | 98.17 ± 0.46 |

**Table 3:** Cell purity assessment by flow cytometry. Table showing percent cells in each fraction stained by each cell-type specific antibody using flow cytometry (n=3, data displayed as average ± standard

| Cell type | O4-PE |  | CD11b-Vioblue |  | CD34-FITC |  | ACSA2-APC |  |
| --- | --- | --- | --- | --- | --- | --- | --- | --- |
|  | Positive (%) | Negative (%) | Positive (%) | Negative (%) | Positive (%) | Negative (%) | Positive (%) | Negative (%) |
| Oligodendrocytes | **99.57 ± 0.12** | 0.29 ± 0.05 | 0.10 ± 0.03 | 99.9 ± 0.00 | 0.70 ± 0.30 | 99.17 ± 0.29 | 3.22 ± 0.62 | 96.60 ± 0.64 |
| Microglia | 3.51 ± 1.13 | 96.27 ± 1.10 | **79.33 ± 2.90** | 20.17 ± 2.78 | 2.37 ± 1.07 | 95.93 ± 1.30 | 2.52 ± 0.61 | 96.10 ± 0.64 |
| Endothelial cells | 14.12 ± 7.86 | 85.17 ± 8.54 | 0.87 ± 0.42 | 99.10 ± 0.43 | **45.77 ± 6.09** | 46.57 ± 5.06 | 12.70 ± 0.55 | 85.80 ± 2.05 |
| Astrocytes | 9.92 ± 3.46 | 89.60 ± 3.34 | 0.68 ± 0.24 | 99.10 ± 0.29 | 29.47 ± 8.51 | 69.17 ± 8.87 | **74.30 ± 0.78** | 23.10 ± 1.26 |
| Neurons | 1.31 ± 0.35 | **98.57 ± 0.34** | 1.25 ± 0.60 | **98.77 ± 0.61** | 3.18 ± 0.18 | **96.43 ± 0.29** | 2.58 ± 0.58 | **97.17 ± 0.54** |

## Quantification and data analysis

RT-qPCR samples were analyzed using the ΔΔCt method with β-actin as the endogenous control gene. In brief, the average Ct of three replicate wells for each sample was subtracted from the β-actin Ct average for an adjusted Ct value (ΔCt). The average Ct of the pre-sorting input samples was subtracted from each sample’s adjusted Ct value (ΔΔCt). The expression fold change was then calculated as 2^−ΔΔCt^ and plotted using GraphPad Prism software (**Figure 2**), and was used to determine the expression levels of each cell type-specific marker in each cell fraction.

The gating strategy for analysis of each cell fraction is affected by its antibody staining and cell size which vary between cell types. For all samples, to determine the population of cells, side scatter (SSC) is plotted against forward scatter (FSC) and the leftmost population of debris is excluded (**Figure 3A, C, E, G, I)**. Next, to select the population of singlets, FSC area (FSC-A) is plotted against FSC height (FSC-H), and doublets are excluded by only selecting the population along the diagonal (singlets) (**Figure 3B, D, F, H, J)**. Next, for each antibody fluorophore, SSC-A is plotted against the fluorophore signal area. This is performed in unstained controls for each cell type and negative/positive gates are set maintaining the positive gate at around <1% for unbiased gating (**Figure 4, pink data points)**. The same gating strategy is applied to stained samples and adjusted as needed to account for variability in staining across different cell types (**Figure 4, black data points)**. Measures of purity and viability are extracted from the plots and listed in table format **(Tables 2 & 3)**.

## Limitations

This protocol isolates oligodendrocytes, microglia, endothelial cells, astrocytes, and neurons from the same mouse brain, establishing a robust foundation for comparative analyses of cellular functions across these cell types. This isolation is performed using pan-cell markers and hence does not select for specific sub-populations of cells. In the case that a specific cell subtype is needed, this protocol can be paired with additional isolation steps to enrich the subtype of interest.

This protocol has been optimized to yield a large number of cells for each cell type and therefore utilizes a large brain region (cerebrum) as input. This results in a loss of spatial resolution, potentially obscuring region-specific cell functions. Further dissection of specific brain regions such as cortex or hippocampus can be performed and used as input in this protocol. Further optimization will be required to determine whether pooling of samples is necessary to achieve a satisfactory cell yield for analysis.

Flow cytometry was used to supplement RT-qPCR to validate the purity of each cell fraction. Flow cytometry proved to be challenging due to the need to compare five cell types of different characteristics using four antibodies of different affinities and brightness. The process of setting gates across different cell types is complex because each cell type has a unique side scatter due to its shape and internal complexity. Furthermore, different side scatter characteristics may be observed in the stained compared to the unstained cells due to the presence of the antibody fluorophores that affect apparent cell morphology. Therefore, the gate for each cell type and fluorophore was set individually. Fluorescence minus one (FMO) controls can be used to try to make improvements in setting the gates for each cell type. However, this will add 20 additional flow samples and prolong the workflow significantly when isolating five cell types characterized by four antibodies. Interchanging the fluorophore tags for each antibody may also help with troubleshooting the gating strategy. Lastly, analyzing each cellular fraction by two cell-specific markers for each cell type might also improve the gating of each cell type.

This protocol utilizes the gentleMACS Octo Dissociator system which is a highly reproducible gentle homogenization system. However, just like any other method of preparation of a cell suspension, this results in the loss of delicate cell processes such as axons and dendrites which may contain specific receptor subtypes and conduct important biological functions like neurotransmission. To understand the full picture of cell-specific roles, this protocol can be paired with other approaches that preserve this level of spatial resolution, such as spatial transcriptomics.

Lastly, this protocol has been tested successfully at ages ranging from 3 to 12 months old. Pilot studies and optimization will be required when using mice aged beyond this range.

## Troubleshooting

### Problem 1

Inadequate enrichment of endothelial cells is observed using flow cytometry staining as compared to RT-qPCR results (Refer to “Quantification and data analysis section”).

### Potential solution

This protocol includes isolation of endothelial cells, which otherwise contaminate the astrocytic and neuronal cell fractions as observed during our optimization phases. A challenge in validating the endothelial cell fraction is that the strong enrichment indicated by RT-qPCR does not match the weak enrichment indicated by flow cytometry. This may be due to the relatively weaker signal of the endothelial cell marker used for flow cytometry compared to other cell type markers. Testing a variety of antibodies from different suppliers with different binding epitopes, fluorescent tags of different wavelengths, and different acquisition parameters may be necessary to achieve a better signal. A combination of two cell marker antibodies may be appropriate. This additional optimization can be pursued if this cell type is of particular interest.

### Problem 2

The homogenized brain suspension is viscous after dissociation (step 5).

### Potential solution

A viscous homogenized brain suspension can be due to over-homogenization of the tissue. This may happen if the brain was cut into pieces that are too small prior to dissociation in the gentleMACS C-tube. The brain should only be cut into pieces roughly 3mm^3^ and not smaller.

### Problem 3

Gradient did not form after layering the resuspension buffer solution with cells over the ovomucoid buffer (step 7).

### Potential solution

A common mistake in step 7 is the use of supplemented EBSS solution instead of the resuspension buffer to resuspend to pelleted cells before layering on the ovomucoid solution. Supplemented EBSS contains trehalose and is denser than the resuspension buffer and will therefore not layer properly on top.

Note: It is normal to see some bigger sized debris clumps deposit to the bottom of the tube (**Figure 1D**). These will be filtered out using a strainer in the following step.

### Problem 4

Lower cell yields are observed.

### Potential solution

Improper handling will impact cell viability. Pipetting and mixing should always be slow and gentle, and the tubes should be kept on ice as much as possible. Incubation with beads and antibodies needs to be in a 4°C fridge and not on ice as this is important for proper binding to the antibody and subsequently better retention on the column. Prewetting strainers and columns is a crucial step to prevent cells getting lodged in the mesh/matrix.

## Resource availability

### Lead contact

Further information and requests for resources and reagents should be directed to and will be fulfilled by the lead contact, Heather Rice.

### Technical contact

Technical questions on executing this protocol should be directed to and will be answered by the technical contact, Heather Rice.

### Materials availability

This study did not generate new unique reagents. All materials used can be purchased from the manufacturers.

### Data and code availability

This paper did not generate any new datasets or code.

## Acknowledgments

This work was supported by the National Institutes of Health (R35GM142726 to H.C.R., R21AG085486 to H.C.R., T32AG052363-08 to S.H., DP5OD033443 to S.R.O., and RF1AG085573 to W.M.F), the Presbyterian Health Foundation (Pilot Research Funding to H.C.R.), the Alzheimer’s Association (SAGA23-1072406 to S.R.O.), and the Veterans Affairs (1I01BX006628 and to W.M.F). Data processing and analysis were supported by the OMRF Center for Biomedical Data Sciences. We thank the OMRF flow cytometry core and members of the Rice, Freeman, Ocañas, and Miller labs for helpful discussions. We thank the representatives from Miltenyi Biotec for their assistance with instrument use and data analysis. The graphical abstract diagrams were created in BioRender. Houmam, S. (2025) https://BioRender.com/kk8aw1j.

## Author contributions

S.H., D.S., S.R.O., W.M.F. and H.C.R. conceived the study. S.H. performed experiments and data analysis. W.M.F. provided access to the MACSQuant10 and Quantstudio5 instruments. K.D.P. and C.S. provided technical assistance. S.H. and H.C.R wrote the manuscript. D.S., C.S., K.D.P, S.R.O., and W.M.F. edited the manuscript.

## References

1. Demmings, M.D., da Silva Chagas, L., Traetta, M.E., Rodrigues, R.S., Acutain, M.F., Barykin, E., Datusalia, A.K., German-Castelan, L., Mattera, V.S., Mazengenya, P., et al. (2024). (Re)building the nervous system: A review of neuron–glia interactions from development to disease. J Neurochem 169, e16258. 10.1111/JNC.16258.

2. Arzalluz-Luqueángeles, and Conesa, A. (2018). Single-cell RNAseq for the study of isoforms— how is that possible? Genome Biol 19, 110. 10.1186/S13059-018-1496-Z.

3. Regan, C., and Preall, J. (2022). Practical Considerations for Single-Cell Genomics. Curr Protoc 2, e498. 10.1002/CPZ1.498.

4. Holt, L.M., Stoyanof, S.T., and Olsen, M.L. Magnetic cell sorting for in vivo and in vitro astrocyte, neuron, and microglia analysis. 10.1002/cpns.71.

5. Schroeter, C.B., Herrmann, A.M., Bock, S., Vogelsang, A., Eichler, S., Albrecht, P., Meuth, S.G., and Ruck, T. (2021). One Brain—All Cells: A Comprehensive Protocol to Isolate All Principal CNS-Resident Cell Types from Brain and Spinal Cord of Adult Healthy and EAE Mice. Cells 10, 651. 10.3390/CELLS10030651.

6. Swartzlander, D.B., Propson, N.E., Roy, E.R., Saito, T., Saido, T., Wang, B., and Zheng, H. (2018). Concurrent cell type–specific isolation and profiling of mouse brains in inflammation and Alzheimer’s disease. JCI Insight 3. 10.1172/JCI.INSIGHT.121109.

7. Ocañas, S.R., Pham, K.D., Blankenship, H.E., Machalinski, A.H., Chucair-Elliott, A.J., and Freeman, W.M. (2022). Minimizing the Ex Vivo Confounds of Cell-Isolation Techniques on Transcriptomic and Translatomic Profiles of Purified Microglia. eNeuro 9. 10.1523/ENEURO.0348-21.2022.

8. Saxena, A., Wagatsuma, A., Noro, Y., Kuji, T., Asaka-Oba, A., Watahiki, A., Gurnot, C., Fagiolini, M., Hensch, T.K., and Carninci, P. (2012). Trehalose-enhanced isolation of neuronal sub-types from adult mouse brain. Biotechniques 52, 381. 10.2144/0000113878.

9. Ramberger, E., Sapozhnikova, V., Ng, Y.L.D., Dolnik, A., Ziehm, M., Popp, O., Sträng, E., Kull, M., Grünschläger, F., Krüger, J., et al. (2024). The proteogenomic landscape of multiple myeloma reveals insights into disease biology and therapeutic opportunities. Nat Cancer 5, 1267. 10.1038/S43018-024-00784-3.

10. Sobue, A., Komine, O., Hara, Y., Endo, F., Mizoguchi, H., Watanabe, S., Murayama, S., Saito, T., Saido, T.C., Sahara, N., et al. (2021). Microglial gene signature reveals loss of homeostatic microglia associated with neurodegeneration of Alzheimer’s disease. Acta Neuropathol Commun 9, 1–17. 10.1186/S40478-020-01099-X/FIGURES/6.

11. Li, Q., Zhu, Z., Wang, L., Lin, Y., Fang, H., Lei, J., Cao, T., Wang, G., and Dang, E. (2021). Single-cell transcriptome profiling reveals vascular endothelial cell heterogeneity in human skin. Theranostics 11, 6461. 10.7150/THNO.54917.

